# Oxamniquine resistance alleles are widespread in Old World *Schistosoma mansoni* and predate drug deployment

**DOI:** 10.1101/657056

**Authors:** Frédéric D. Chevalier, Winka Le Clec’h, Marina McDew-White, Vinay Menon, Meghan A. Guzman, Stephen P. Holloway, Xiaohang Cao, Alexander B. Taylor, Safari Kinung’hi, Anouk N. Gouvras, Bonnie L. Webster, Joanne P. Webster, Aidan M. Emery, David Rollinson, Amadou Garba Djirmay, Khalid M. Al Mashikhi, Salem Al Yafae, Mohamed A. Idris, Hélène Moné, Gabriel Mouahid, P. John Hart, Philip T. LoVerde, Timothy JC. Anderson

**Affiliations:** Texas Biomedical Research Institute, San Antonio, Texas, USA; Departments of Pathology University of Texas Health Science Center at San Antonio, San Antonio, Texas, USA; Biochemistry & Structural Biology, University of Texas Health Science Center at San Antonio, San Antonio, Texas, USA; X-ray Crystallography Core Laboratory, University of Texas Health Science Center at San Antonio, San Antonio, Texas, USA; National Institute for Medical Research, Mwanza, United Republic of Tanzania; London Centre for Neglected Tropical Disease Research (LCNDTR), Imperial Collge, London, United Kingdom; Wolfson Wellcome Biomedical Laboratories, Natural History Museum, London, United Kingdom; Centre for Emerging, Endemic and Exotic Diseases (CEEED), Royal Veterinary College, University of London, United Kingdom; Réseau International Schistosomiases Environnemental Aménagement et Lutte (RISEAL), Niamey, Niger; World Health Organization, Geneva, Switzerland; Directorate General of Health Services, Dhofar Governorate, Salalah, Sultanate of Oman; Sultan Qaboos University, Muscat, Sultanate of Oman; Host-Pathogen-Environment Interactions laboratory, University of Perpignan, France

## Abstract

Do mutations required for adaptation occur *de novo*, or are they segregating within populations as standing genetic variation? This question is key to understanding adaptive change in nature, and has important practical consequences for the evolution of drug resistance. We provide evidence that alleles conferring resistance to oxamniquine (OXA), an antischistosomal drug, are widespread in natural parasite populations under minimal drug pressure and predate OXA deployment. OXA has been used since the 1970s to treat *Schistosoma mansoni* infections in the New World where *S. mansoni* established during the slave trade. Recessive loss-of-function mutations within a parasite sulfotransferase (SmSULT-OR) underlie resistance, and several verified resistance mutations, including a deletion (p.E142del), have been identified in the New World. Here we investigate sequence variation in *SmSULT-OR* in *S. mansoni* from the Old World, where OXA has seen minimal usage. We sequenced exomes of 204 *S. mansoni* parasites from West Africa, East Africa and the Middle East, and scored variants in *SmSULT-OR* and flanking regions. We identified 39 non-synonymous SNPs, 4 deletions, 1 duplication and 1 premature stop codon in the *SmSULT-OR* coding sequence, including one confirmed resistance deletion (p.E142del). We expressed recombinant proteins and used an *in vitro* OXA activation assay to functionally validate the OXA-resistance phenotype for four predicted OXA-resistance mutations. Three aspects of the data are of particular interest: (i) segregating OXA-resistance alleles are widespread in Old World populations (4.29 – 14.91% frequency), despite minimal OXA usage, (ii) two OXA-resistance mutations (p.W120R, p.N171IfsX28) are particularly common (>5%) in East African and Middle-Eastern populations, (iii) the p.E142del allele has identical flanking SNPs in both West Africa and Puerto Rico, suggesting that parasites bearing this allele colonized the New World during the slave trade and therefore predate OXA deployment. We conclude that standing variation for OXA resistance is widespread in *S. mansoni*.

**AUTHOR SUMMARY:** It has been argued that drug resistance is unlikely to spread rapidly in helminth parasites infecting humans. This is based, at least in part, on the premise that resistance mutations are rare or absent within populations prior to treatment, and take a long time to reach appreciable frequencies because helminth parasite generation time is long. This argument is critically dependent on the starting frequency of resistance alleles – if high levels of “standing variation” for resistance are present prior to deployment of treatment, resistance may spread rapidly. We examined frequencies of oxamniquine resistance alleles present in *Schistosoma mansoni* from Africa and the Middle East where oxamniquine has seen minimal use. We found that oxamniquine resistance alleles are widespread in the Old World, ranging from 4.29% in the Middle East to 14.91% in East African parasite populations. Furthermore, we show that resistance alleles from West African and the Caribbean schistosomes share a common origin, suggesting that these alleles travelled to the New World with *S. mansoni* during the transatlantic slave trade. Together, these results demonstrate extensive standing variation for oxamniquine resistance. Our results have important implications for both drug treatment policies and drug development efforts, and demonstrate the power of molecular surveillance approaches for guiding helminth control.

## INTRODUCTION

The rate at which drug resistance alleles (or any other beneficial alleles) spread within populations in response to a drug treatment (or any other selection pressure) is critically dependent on the starting frequency of resistance alleles in the population when new drugs are deployed (1, 2). If no such alleles are present when a novel drug is introduced, then there is a waiting time for resistance alleles to arise. Furthermore, because the starting allele frequency will be 1/(2N_e_), where N_e_ is the effective population size, the vast majority of resistance alleles that arise will be lost due to genetic drift and fail to establish (2, 3) (Fig. 1). The barrier to establishment is particularly severe for recessive traits because these will be present in heterozygotes at low frequency and therefore not exposed to selection. This effect, otherwise known as “Haldanes’s sieve” (4), may be particularly relevant for drug resistance evolution in diploid pathogens, because resistance mutations in drug targets typically result in phenotypic resistance only when two copies are present. However, if resistance alleles are already segregating as standing variation in pathogen populations when drug treatment is initiated, then there is no waiting time for mutations to arise. Furthermore, there is a high probability of fixation and spread is rapid because many resistance alleles are present in homozygous recessive genotypes and therefore exposed to selection. Hence, whether resistance alleles (or alleles for other traits) arise *de novo* or are segregating as standing variation, is a central question in evolutionary biology and public health, and key to predicting the effective “shelf life” of new drug treatments (3, 5).

**Fig. 1.**
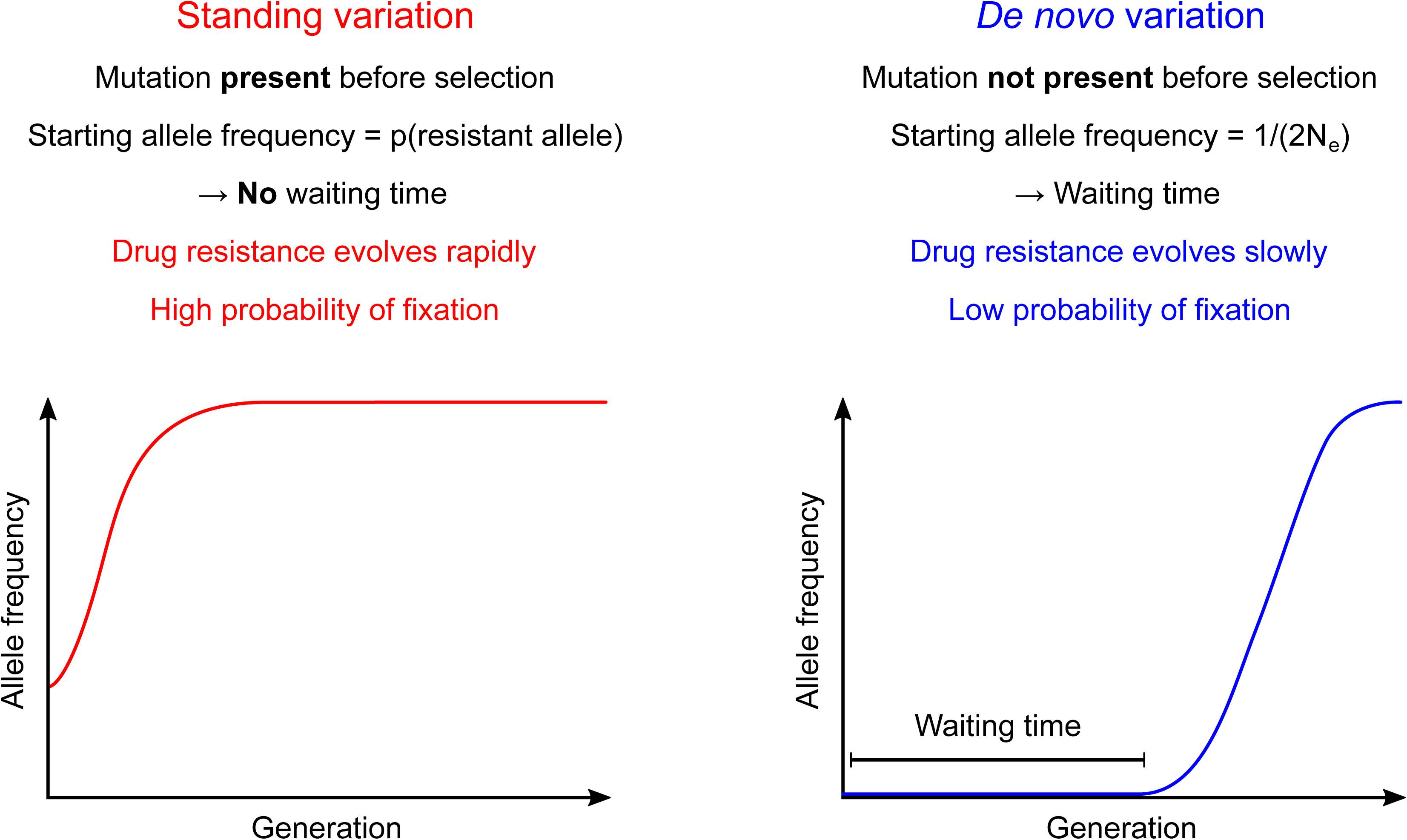
Drug resistance evolution from standing variation or *de novo* mutation. Drug resistance alleles spread rapidly and reach fixation with a high probability when drug resistance alleles are already present as standing variation. In contrast, if resistance alleles are absent when treatment is initiated, resistance mutation must arise *de novo*, so there is a waiting time before the resistance alleles appear, and most resistance alleles are lost by genetic drift, so the probability of establishment and fixation is low.

There is growing evidence for resistance to anti-helminthics in several helminths infecting humans, including *Onchocerca volvulus* (6), the filarial nematode causing river blindness, and in soil transmitted helminths (hookworm, whipworm and Ascaris roundworms) (7) which cumulatively infect over one billion people worldwide (8). However, perhaps the best understood is oxamniquine resistance in schistosome blood flukes, for which the mechanism of drug action and the genetic and molecular basis for drug resistance are now known. Oxamniquine (OXA) kills Schistosoma mansoni, but is ineffective against two other major schistosome species infecting humans (*S. haematobium* and *S. japonicum*) (9). OXA is a pro-drug that is activated by a schistosome sulfotransferase (*SmSULT-OR*) encoded on chromosome 6 (10). Loss-of-function mutations within *SmSULT-OR* result in resistance: only parasites that are homozygous for resistance alleles are phenotypically resistant (OXA-R). We initially identified the locus and mutation, a single amino acid deletion (p.E142del), underlying OXA-R using a genetic cross involving a resistant parasite line selected from a parasite isolate from a Puerto Rican patient in 1971 (11). Additionally we identified a second mutation (p.C35R) in *S. mansoni* obtained from an incurable Brazilian patient (12). These two mutations result in disruption of the active site of the *SmSULT-OR* and inability to activate OXA (10). Both p.E142del and p.C35R, and two additional mutations resulting in truncated proteins, were subsequently identified in a field survey of *SmSULT-OR* genetic variation of parasites from a single Brazilian village (13). The combined allele frequency of these four resistant alleles was 1.85%.

OXA was widely used in the New World, where only *S. mansoni* is present, during the 1970s to early 2000’s (14), so the OXA-R alleles we observed in Brazil (13) could conceivably have resulted from drug selection. In contrast, OXA saw minimal usage in Africa where both *S. haematobium* and *S. mansoni* are present. In South America, an estimated 6 million doses of OXA were used to treat the 7 million people infected with *S. mansoni* prior to 1987 (15). In contrast, 3 million doses had been used in Africa (Egypt, Ethiopia, Ivory Coast, Kenya, Malagasy, Malawi, Rwanda-Burundi, South Africa, Sudan, Tanzania, Uganda, Zambia, Zaire) and the Middle East (Arabian penensula) (15), where an estimated ∼60 million people were infected with *S. mansoni* (16). Hence, OXA selection was minimal in Africa compared with South America. The central aim of this paper is to determine whether *de novo* mutation or standing variation best explains OXA-R in *S. mansoni* populations. To do this we examined *SmSULT-OR* genetic variation in Old World parasite populations where OXA treatment has been minimal.

## RESULTS

### 1. Samples

We sequenced exomes from 92 miracidia from Tanzania (n=57), Niger (n=10), and Senegal (n=25) from the SCAN collection at the Natural History Museum (17), with each miracidium derived from a different patient. In addition, we included 112 samples collected from Oman: these included 86 single worms and 26 pools of cercariae from single infected snails. The Omani samples were derived from 11 human infections, 2 naturally infected rodents, and 26 naturally infected snails (Table 1). We obtained an average of 15.7 ± 8 (mean ± standard deviation, s.d.) million reads per library (supp. table 1). A very high proportion of these reads were mapped to the reference genome (91.9 ± 6.29 (s.d.) % on average). We captured on average 99.08 ± 3.92 (s.d.) % of the bait regions. We obtained an average read depth of 64.40 ± 32.27 (s.d.) in the bait regions. For *SmSULT-OR*, we obtained sequence across 95.06 ± 6.54 (s.d.) % of the coding sequence with an average read depth of 43.09 ± 26.96 (s.d.).

**Table 1.** Summary statistics for sequence variation in *SmSULT-OR*.

| Location | Sample size | Parasite stage sampled | Host | Sequence | Notes<sup>b</sup> |
| --- | --- | --- | --- | --- | --- |
| South America (Brazil)<sup>a</sup> | 189 | Miracidia | Human | Sanger sequence of *SmSULT-OR* exons | 1-11 miracidia from each of 49 people from single village (13) |
| West Africa (Niger/Senegal) | 25/10 | Miracidia | Human | Exome sequence | 1 miracidia from each person |
| East Africa (Tanzania) | 57 | Miracidia | Human | Exome sequence | 1 miracidia from each person |
| Middle East (Oman) | 112 | Miracidia/adult worms/cercariae | Human/Rat/Snail | Exome sequence | 1-53 parasite genotypes analysed per host from 11 people, 2 rats and 26 snails from 8 locations |
| Caribbean (Puerto Rico)<sup>a</sup> | 1 | Adult worms | Hamster | Genome sequence | OXA-resistant parasite used in a genetic cross (10) |

<sup>a</sup> published data836 <sup>b</sup> see supplementary table 2 for detailed information on sample locations

We compared these Old World parasite samples with: (i) the OXA-R parasite from Puerto Rico (HR) carrying the causative p.E142del mutation identified previously (10), for which whole genome sequence is available, and (ii) published *SmSULT-OR* sequences from a single Brazilian location (n=189) (13) (Table 1).

### 2. New and known mutations identified in the *SmSULT-OR* gene

We identified a total of 85 *SmSULT-OR* mutations across all three populations (West Africa, East Africa and Middle East), including 76 coding mutations and 9 non-coding mutations (supp. table 2). Exon 1 carried 24 mutations. These included 13 non-synonymous single nucleotide polymorphisms (SNPs), 8 synonymous SNPs, one duplication, one deletion, and one premature stop codon. Exon 2 carried 52 mutations: 26 non-synonymous SNPs, 23 synonymous and three deletions. There were no differences in the density of mutations in the two exons (Fisher’s exact test, d.f.=1, p=0.09). The overall transition/transversion ratio was 1.94. Forty-three percent (37/85 mutations) showed an average allele frequency greater than 5%.

**Table 2.**
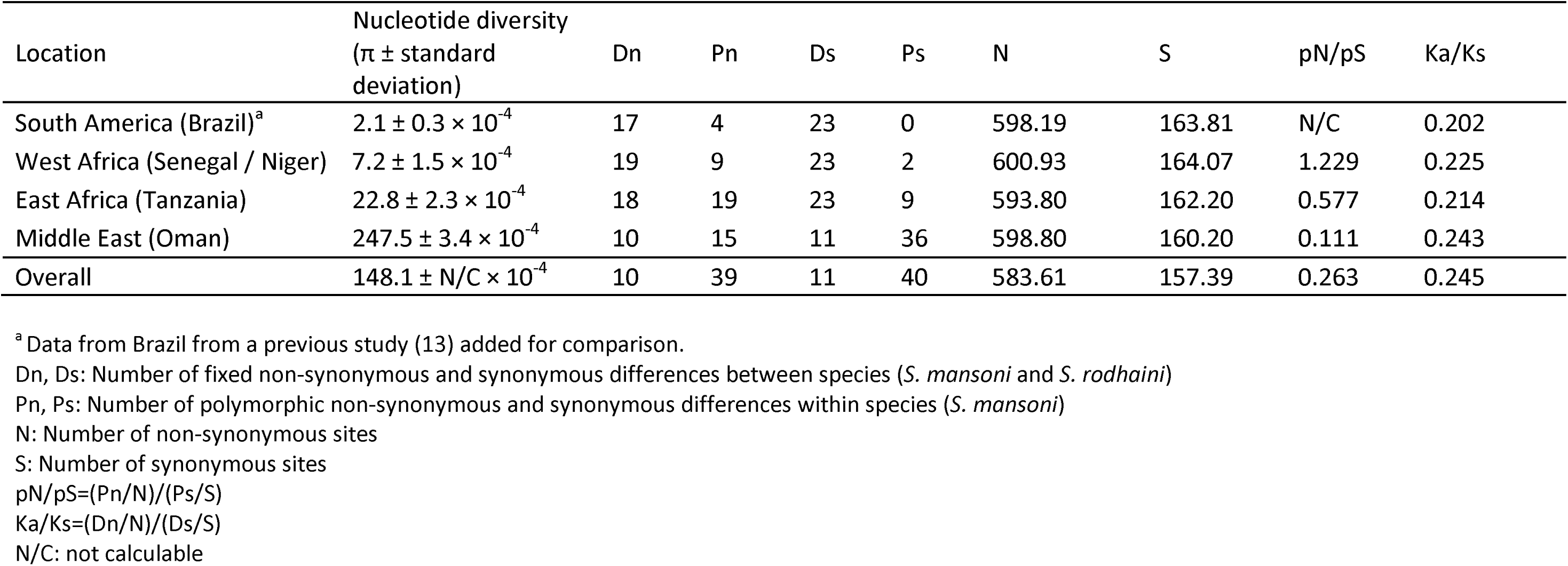
Summary statistics for sequence variation in *SmSULT-OR*.

Among the 85 mutations, we have previously found three coding and one non-coding mutations in the New World (Caribbean and Brazil): the substitution p.P67L (g.200C>T), the deletion p.E142del (g.4348_4350delGAA), the substitution p.L256W (g.4691T>G) and the 3’ untranslated region (UTR) substitution g.4720C>T (10, 13). Just one of these mutations (p.E142del) encodes a confirmed OXA-R allele. Frequencies of all mutations in each population are shown in supp. table 2.

We measured nucleotide diversity (π) in both New and Old World populations. Nucleotide diversity was the highest in Oman (π = 0.02424 ± 2.4 × 10^−4^) followed by East Africa (π = 0.00195 ± 2 × 10^−4^), West Africa (π = 0.00126 ± 1.7 × 10^−4^) and Brazil (π = 0.00021 ± 0.3 × 10^−4^). If drug pressure drives the changes at *SmSULT-OR*, we might expect to see signals of positive selection. We examined sequence conservation by measuring proportions of non-synonymous difference per non-synonymous site relative to synonymous changes per synonymous site (K_a_/K_s_) between *S. mansoni* and a *S. rodhaini* outgroup. K_a_/K_s_ ranged from 0.20 – 0.24 suggesting that changes at non-synonymous sites are 4-5 times less abundant than at synonymous sites, consistent with weak purifying selection. Similarly, the ratio of non-synonymous polymorphisms per non-synonymous site relative to synonymous polymorphisms per synonymous site (pN/pS) ranged from 0.111 to 1.36 and none of these values were significantly greater than unity (Table 2). These tests did not provide evidence that *SmSULT-OR* is under positive selection.

### 3. *in vitro* tests and in silico evaluation of impact of mutations

We used three approaches to identify *SmSULT-OR* mutations that are likely to result in OXA-R.

#### a. Visual assessment

Some mutations have obvious deleterious effect on the protein by introducing premature stop codons. This is the case for three mutations identified in *SmSULT-OR*. These mutations will generate truncated proteins that do not contain the active site. Other mutations, such as p.L179P, p.P225S or p.W120R, were either close enough to the OXA or the 3’-phosphoadenosine-5’-phosphosulfate (PAPS) binding sites or showed potential steric hindrance or structural deformation likely to have an effect on protein activity. Other mutations, such as p.A74T, p.P106S or p.Q176R, were on the surface of the protein, distant to any active site, and therefore unlikely to have an impact.

#### b. Functional assay of OXA binding

We used an *in vitro* OXA activation assay to functionally assay the impact of mutations on *SmSULT-OR* activity for four newly discovered mutations and the two known OXA-R mutations (Fig. 3). The known resistant mutations, p.C35R and p.E142del, showed from zero to very low level of OXA activation, as expected (10). p.L179P showed a similarly low level of activation. The mutations p.S160L and p.P225S showed intermediate OXA activation: we conservatively classed these as OXA sensitive alleles. p.P106S showed similar activity to the wild type and p.S160L. We were able to produce recombinant protein p.W120R but all our attempts to fold this protein were unsuccessful. This mutation clearly has a dramatic impact on protein stability as predicted by visual inspection and thermodynamic modelling results (below). We considered this mutation to be OXA-R.

**Fig. 2.**
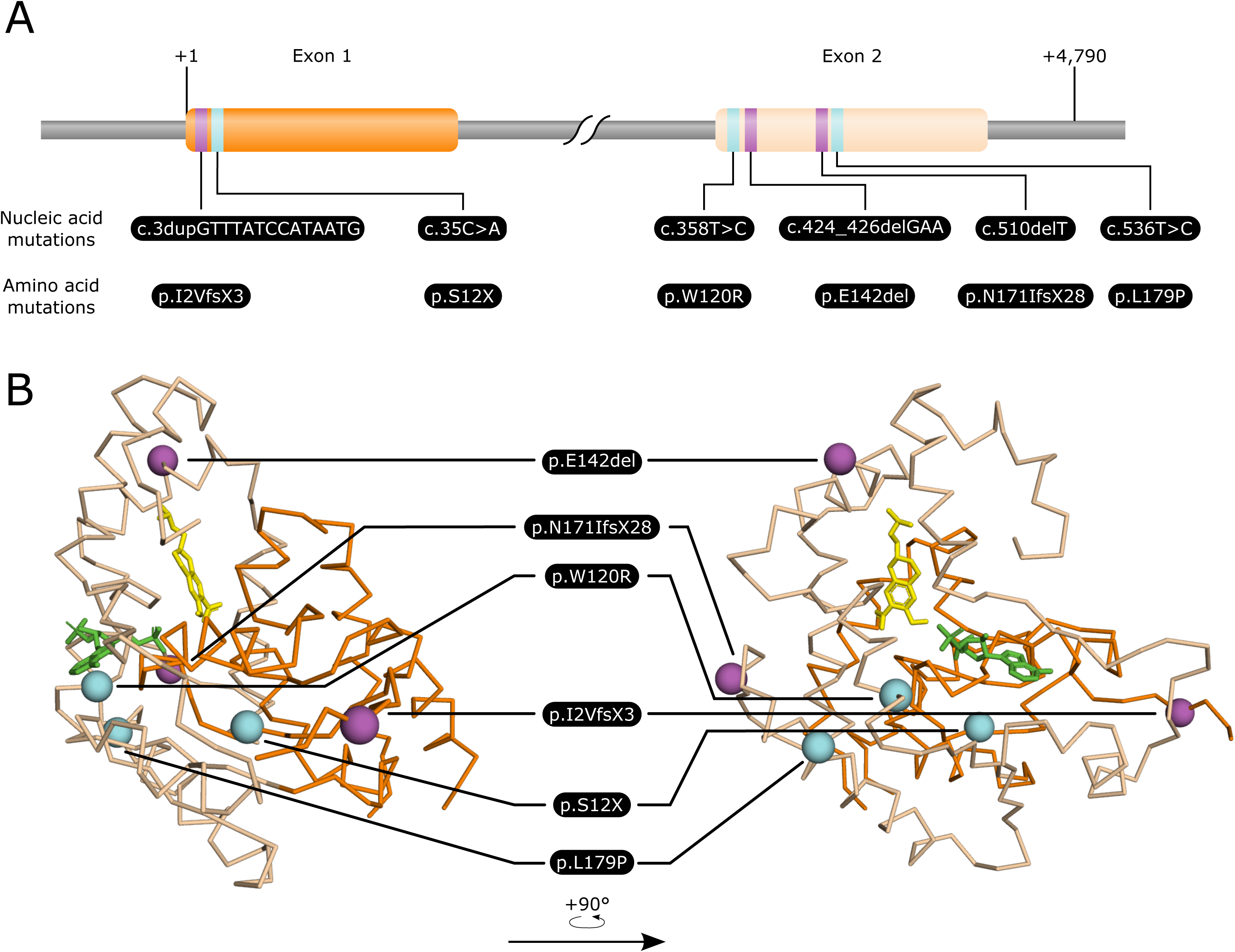
Mapping of the resistance mutations on the gene sequence and structure of *Schistosoma mansoni* SmSULT-OR sulfotransferase. Exon 1 and exon 2 are represented in orange and beige, respectively. Single nucleotide polymorphisms and insertion/deletion events are represented in cyan and magenta, respectively. (A) Linear representation of the *SmSULT-OR* gene showing the relative position of the mutations and their translation in amino acid sequences. (B) Positions of mutations on the *SmSULT-OR* protein. Oxamniquine is represented in yellow, 3’-phosphoadenosine-5’-phosphosulfate (PAPS) co-factor is represented in green.

**Fig. 3.**
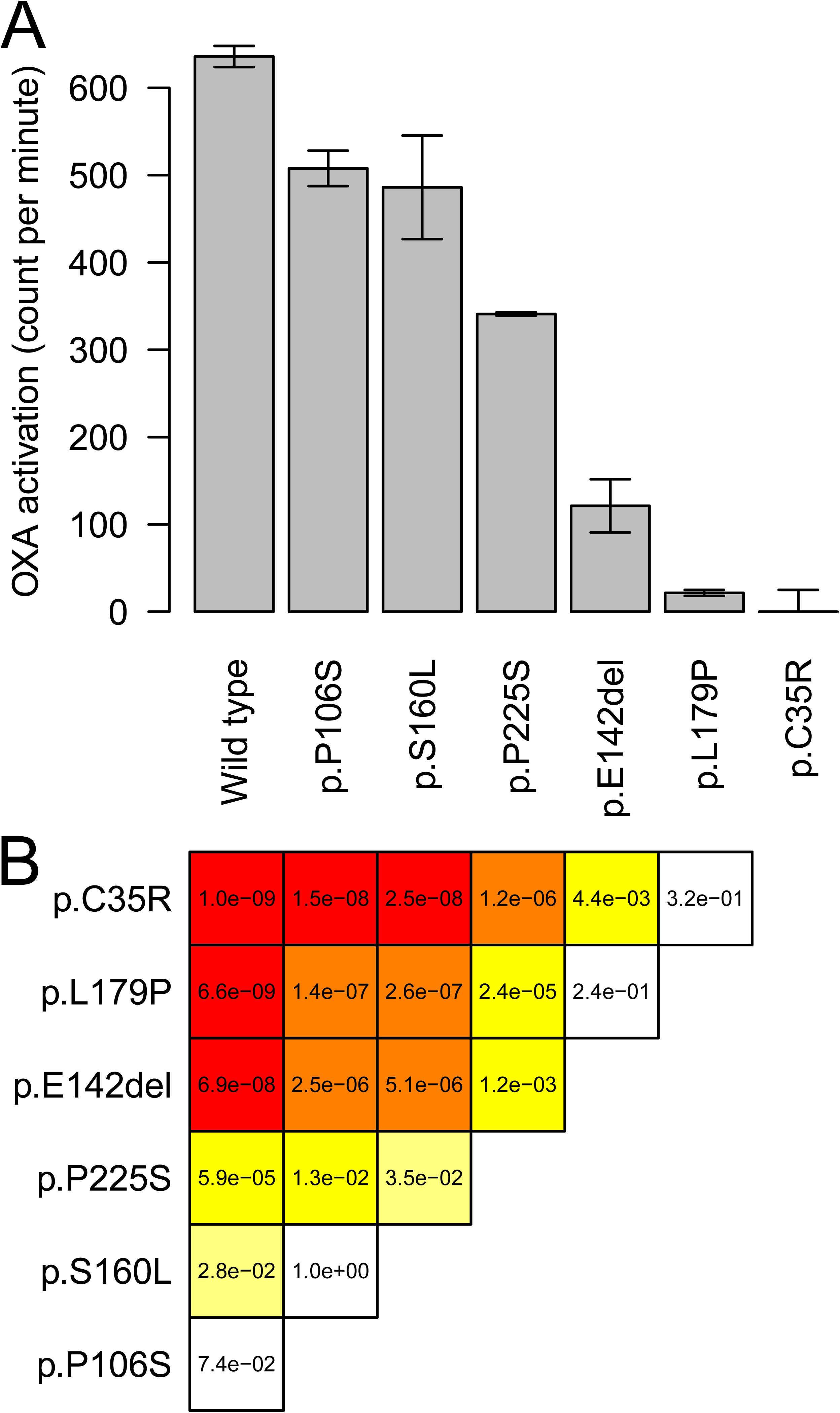
Enzymatic activity of recombinant *Schistosoma mansoni* SmSULT-OR sulfotransferase expressed from different allelic variants. This *in vitro* oxamniquine activation assay quantifies DNA-oxamniquine complexes by scintillation (counts per minute). (A) Bars show the mean of three replicates, while error bars are S.E.M. (B) The triangular matrix shows the p-values from the pairwise comparisons (Tukey’s HSD, p < 0.05). The color is proportional to the level of significance, from white (not significant) to red (most significant). Enzyme carrying known loss-of-function mutations, such as p.C35R or p.E142del, as well as a newly identified variant (p.L179P) showed no or low oxamniquine activation, while two newly identified variants (p.S160L and p.P225S) showed intermediate activation. The newly identified p.P106S did not impair oxamniquine activation.

#### c. Thermodynamic modelling

We used thermodynamic modelling to evaluate the potential impact of substitutions on protein stability (indels such as p.E142del cannot be modelled). The difference in free enthalpy (*ΔΔG*) ranged from −2.942 to 19.826 for the 37 mutations (Fig. 4). Among those, p.L179P and p.C35R which have the greatest impact on OXA activation (Fig. 3) showed the second and third highest *ΔΔG*, respectively. p.W120R, for which we were unable to fold recombinant protein, showed the highest *ΔΔG*, consistent with a dramatic impact on stability. p.S160L and p.P225S showed intermediate *ΔΔG* consistent with the results of the OXA activation assay.

**Fig. 4.**
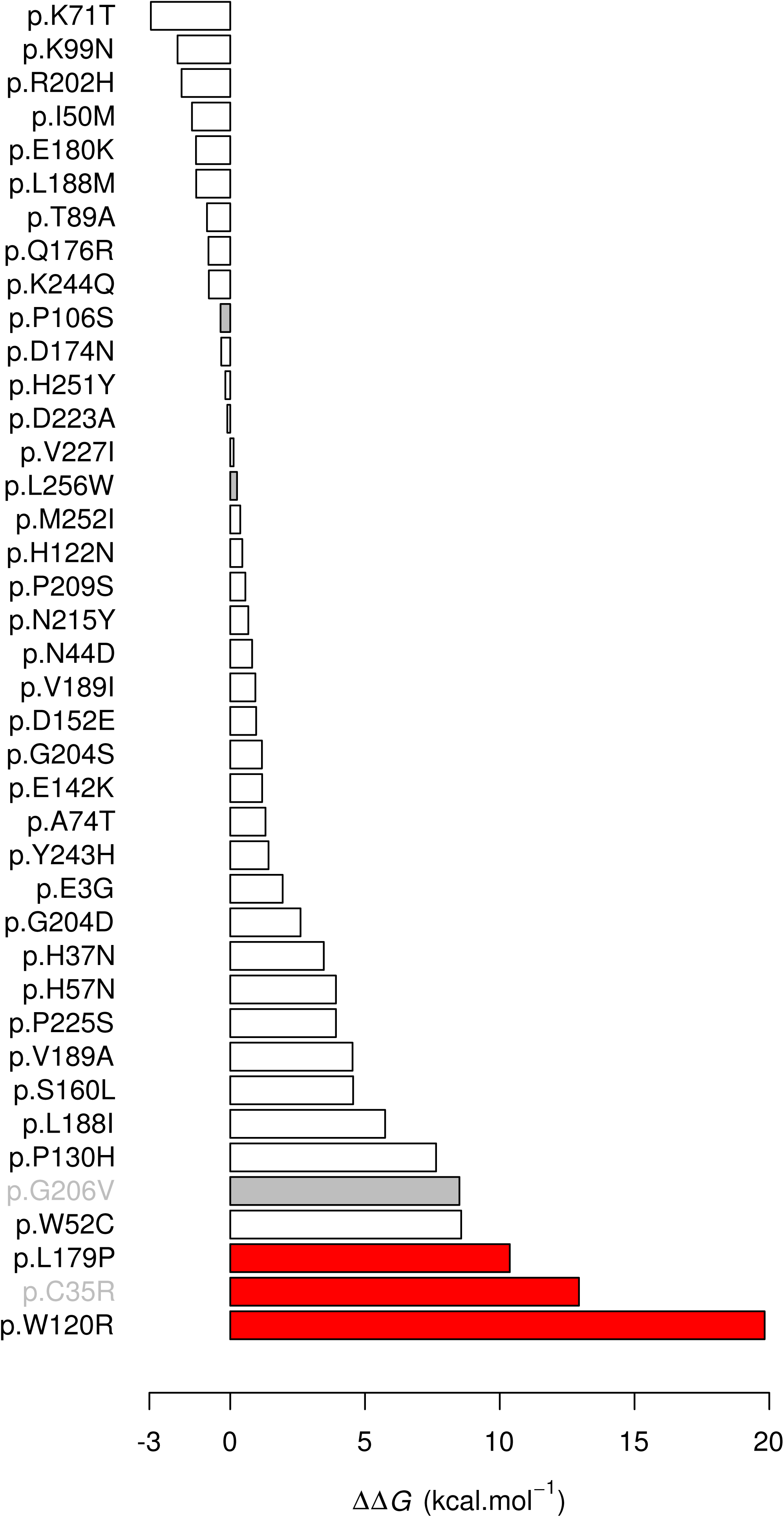
In silico evaluation of mutations on protein stability. We computed the difference in free enthalpy (*ΔΔG*) between the mutated and the wild type proteins. The higher the *ΔΔG*, the more unstable is the mutated protein. Only single amino acid changes within the resolved crystal structure were examined. For completeness, we included mutations from our current dataset and from previous studies (10, 13). Grey bars correspond to known sensitive alleles. Red bars correspond to validated resistant alleles. Grey labels correspond to mutations identified previously from South America (13).

In total, we identified 7 independent OXA-R mutations in Old and New World parasite populations examined using these three approaches (Table 3, Fig. 2). These included 3 indels and 4 amino acid mutations.

**Table 3.** OXA-R mutations scored in the *Schistosoma mansoni SmSULT-OR* gene and their frequency in the New and Old World.

| Nucleic acid mutation | Amino acid mutation | Mutation frequency |  |  |  |
| --- | --- | --- | --- | --- | --- |
|  |  | South America (Brazil)<sup>a</sup><br>n=189 | West Africa (Senegal / Niger)<br>n=25 / n=10 | East Africa (Tanzania)<br>n=57 | Middle East (Oman)<br>n=112 |
| c.3dupGTTTATCCATAATG | p.I2VfsX3 | - | - | 0.0096 | - |
| c.35C>A | p.S12X | - | 0.014 | - | - |
| c.103T>C | p.C35R | 0.0027 | - | - | - |
| c.358T>C | p.W120R | - | - | 0.0510 | 0.065 |
| c.424_426delGAA | p.E142del | 0.008 | 0.035 | 0.0098 | - |
| c.510delT | p.N171IfsX28 | - | - | 0.0625 | - |
| c.536T>C | p.L179P | - | - | 0.0089 | - |
| Total frequency |  | 0.0107 | 0.044 | 0.1418 | 0.065 |

The code used for nucleic acid mutations indicates the sequence type (c = coding), the position, and the mutation type (X>Y = substitution of X by Y, insN = insertion of N, delN = deletion of N, dup = duplication). The code used for amino acid mutations indicates the sequence type (p = protein), the reference amino acid, the position, and finally the alternative amino acid, and when frame shift (fs) occurs, the position of the stop codon (X) after the mutation. For details about the nomenclature, see Ogino *et al.* (79).
<sup>a</sup> Data from Brazil were obtained in a previous study (13) and added for comparison.

### 4. Frequency of OXA-R alleles

Having determined which of the mutations identified are likely to cause OXA-R, we measured the frequency of resistant variants and the frequency of resistant parasites in each population (Table 3, Fig. 5). To accurately identify resistant alleles and parasites, we phased our variant calling data on the first 3 Mb of the chromosome 6 (corresponding to 30,812 variable sites). West African samples carried only two resistant variants (p.S12X and p.E142del). East African samples carried five resistant variants, two of them at a frequency over 0.05 (p.W120R and p.N171IfsX28). The p.W120R mutation was also found at high frequency in Middle East (0.065). We found the highest frequency of OXA-R alleles in East Africa (14.91%, 17/114), followed by the Middle East (6.25%, 14/224), West Africa (4.29%, 3/70), and Brazil (1.85%, 7/378).

**Fig. 5.**
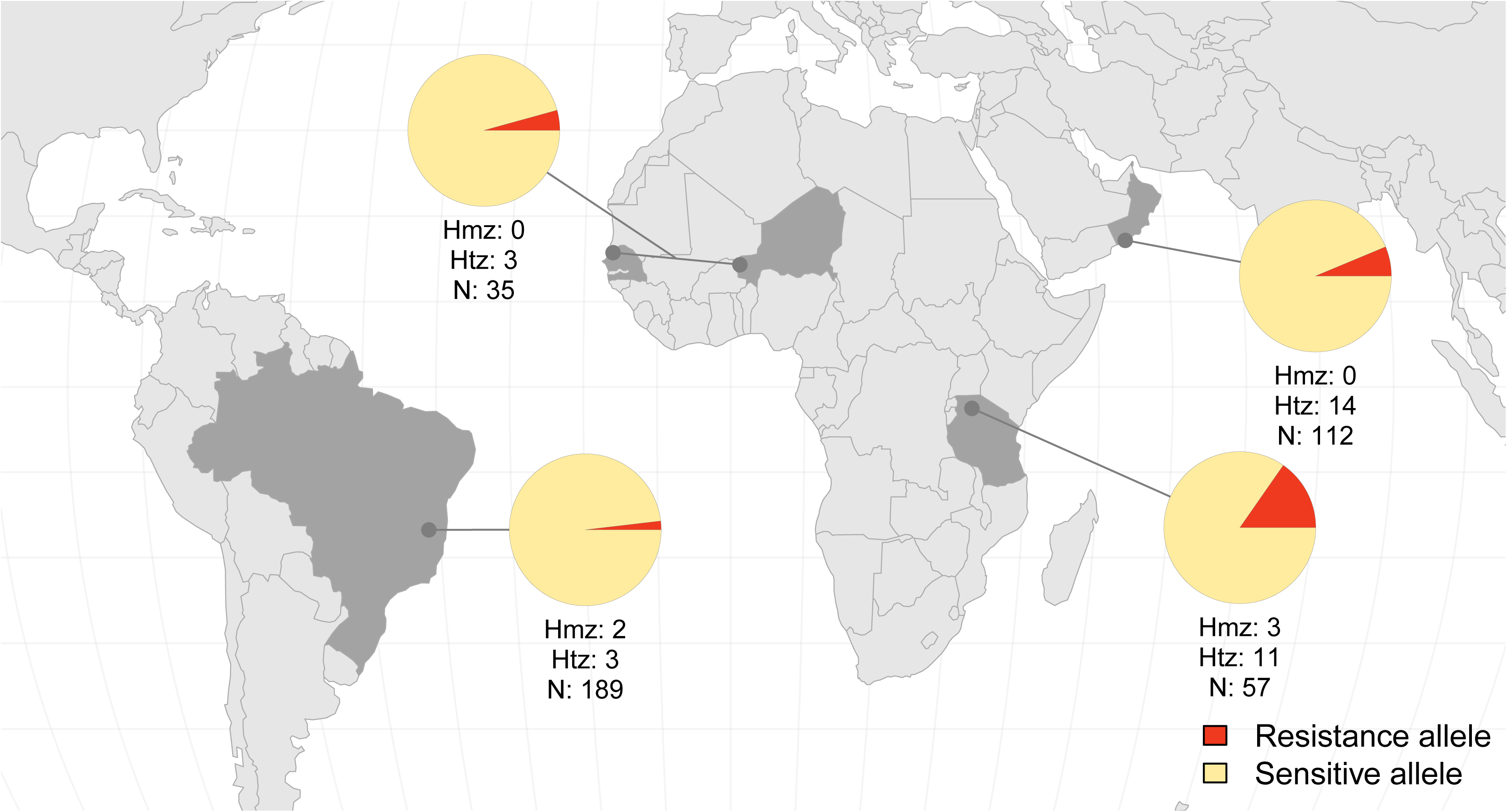
World map showing proportion of resistance and sensitive alleles in South America (Brazil), West Africa (Senegal and Niger), East Africa (Tanzania) and Middle East (Oman). Number of homozygous (hmz) and heterozygous (htz) parasites for resistant alleles, and total number of parasites sampled (n) are shown below the pie charts. East Africa showed the highest frequency of resistant alleles and resistant parasites. Data from Brazil from a previous study (13) was added for comparison.

Most parasites carried OXA-R alleles in the heterozygous state and are predicted to be OXA sensitive. Heterozygous OXA-sensitive parasites were found in West and East Africa (8.6% and 24.6%, respectively), Middle East (13.4%) and in Brazil (1.06%). Parasites that are homozygous for OXA-R alleles at *SmSULT-OR* (but not necessarily carrying the same resistant variant) are OXA-R. We found homozygous parasites predicted to be phenotypically OXA-R at a frequency of 5.26% (3/57) in East Africa and 1.06% (2/189) in South America only (Fig. 5).

### 5. Haplotype analysis of *SmSULT-OR* and flanking regions

We used our phased data to investigate the haplotypes surrounding our resistant variant (p.E142del) in the Old and New World (Caribbean) (Table 1) and to investigate whether these resistant alleles derived from a common ancestor. This is of interest because it would suggest that these alleles predate the slave trade. We identified an identical 102.5 kb haplotype block around p.E142del shared between the Caribbean sample and one of the West African samples from Niger (Fig. 6A). This haplotype block contained 399 SNPs that varied in at least one of the 205 parasites examined (all samples except those from Brazil where haplotype data were not available). We used a minimum spanning network to investigate haplotype relationships using the 399 bi-allelic variants found in this 102.5 kb block across all phased haplotypes (Fig. 6B). The haplotypes associated with the p.E142del OXA-R alleles found in both Puerto Rico and Niger clustered together, consistent with a common origin for West African and Caribbean p.E142del OXA-R mutations.

**Fig. 6.**
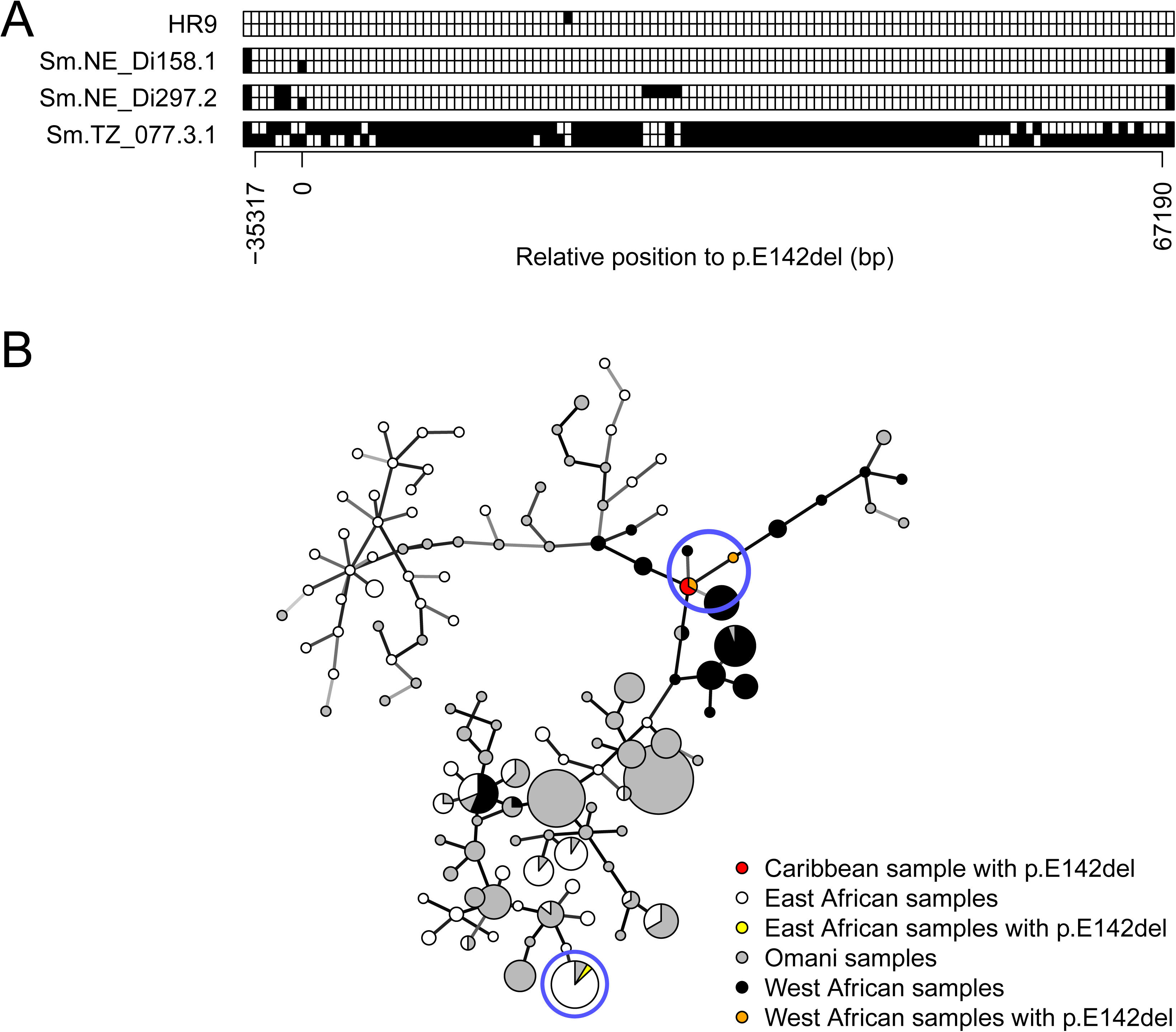
Common origin of the p.E142del mutation in the Old and New World. (A) Haplotype variation in all the samples bearing p.E142del from Caribbean (HR9), Niger (NE) and Tanzania (TZ) across a 102.5 kb region of chr. 6. Each row represents a chromosome, HR9 was used as reference, blank squares reflect HR9 allele state, and black squares correspond to the alternative allele. Relative bp position to p.E142del (0 bp) is shown on the x-axis. The first and last variants showed on the block correspond to the break of the haplotype block. The Caribbean sample (HR9) and a Nigerien sample (Sm.NE_Di158.1) share an identical haplotype block of 102.5 kb. (B) Minimum spanning network of 410 haplotypes of the 102.5 kb region previously identified. The network was built using 399 bi-allelic variants. Node size is proportional to sample size (smallest node: n=1; biggest node n=50). Nodes with samples carrying p.E142del are circled in blue. Caribbean and West African haplotypes carrying p.E142del clustered together indicating a common origin. The p.E142del from East Africa has different flanking haplotypes.

The p.E142del mutation was also identified in Brazil (13) but we sequenced the *SmSULT-OR* exons only, rather than complete exomes, for these Brazilian samples preventing reconstruction of extended haplotypes. This Brazilian p.E142del was also found associated with two mutations (p.P67L and g.4720C>T) found in West Africa but without the p.L256W found in samples from the Caribbean and Niger. Similarly, the p.E142del was also found in a Tanzanian sample but this sample has a distinct haplotype from those found in West Africa (Fig. 6). These results suggest either that there are several independent origins of East and West African p.E142del mutations, or alternatively, that the p.E142del is extremely old and has distinctive flanking haplotypes resulting from recombination and mutation.

## DISCUSSION

### 1. Evidence for standing genetic variation for OXA resistance

Two lines of evidence support the view that standing variation is the source of OXA-R alleles in schistosomes. First, we have indirect evidence: OXA-R alleles are geographically widespread in Africa and the Middle East despite limited treatment with OXA (or hycanthone, its structural analog; (18)) in these regions. Furthermore, that OXA-R alleles are found at high frequency in East Africa, where there has been minimal use of OXA, while OXA-R alleles are found at low frequency in Brazil, where OXA has been used extensively, is also consistent with the idea that mutation and drift, rather than selection, explain the patterns of variation observed. The second line of evidence is more direct: we showed that an OXA-R mutation, p.E142del, with an identical 102.5 kb flanking haplotype, is sampled from both West African and Caribbean schistosomes. This result strongly suggests that p.E142del alleles were present in West Africa prior to the transatlantic slave trade (1501-1867) (19), and were transferred to the New World on ships carrying West African slaves. Hence, OXA-R alleles were segregating within *S. mansoni* populations at least 470 years before deployment of OXA in the 1970s (20).

Our molecular data are backed up by clinical observations from the early use of OXA. Resistant parasites were detected in Brazil in the 1970s (21, 22), before any mass drug administration (14). Similar observations of *S. mansoni* infected patients resistant to OXA treatment were made in East Africa (23). The existence of OXA-R parasites was confirmed experimentally by infecting mice with parasites isolated from patients that were parasite positive following drug treatment (21–23). Together these observations support the existence of segregating OXA-R alleles in schistosome populations before OXA deployment for parasite control.

### 2. Differences in OXA treatment efficacy between East and West Africa

Given the variation in OXA-R frequencies that we observe in different geographical samples, it is interesting to examine the literature on clinical treatment efficacy of OXA for treating schistosomiasis patients. Early trials of OXA to treat patients with intestinal schistosomiasis resulted in multiple treatment failures in Egypt, East Africa and South Africa (23–27). As a consequence, the WHO recommended use of higher doses of OXA in East Africa, compared with West Africa (28). Human host metabolism did not explain this lack of efficacy because the OXA availability in blood was the same or higher in the East African patient populations than in West African or South American populations (29, 30). Therefore, the presence of resistant parasites was the most likely explanation. OXA-R parasites were identified in Kenya by treating mice infected with parasites from patients showing poor treatment response (23). We observed a 14.91% frequency of OXA-R alleles (and 5.26% frequency of homozygous OXA-R parasites) in East Africa compared with 4.29% frequency of OXA-R alleles (and no homozygous OXA-R parasites) in West Africa. Our results provide a molecular explanation for the poor treatment response observed in East Africa, relative to other locations.

### 3. Implications for development of OXA derivatives

OXA kills *S. mansoni* but not S. haematobium or S. japonicum. OXA derivatives are currently under development to obtain more potent molecules acting against all three schistosome species. In fact, OXA derivatives that kill adult worms of the three main species of schistosomes infecting humans *in vitro* have now been developed (31). The low frequency of homozygous OXA-R parasites in *S. mansoni* suggests that OXA derivatives will result in imperfect cure rate for this species, particularly in East Africa. However, the primary purpose of generating OXA derivatives is to treat *S. haematobium* and *S. japonicum*, perhaps as a partner drug for praziquantel (PZQ). Examining natural variation in homologous sulfotransferase genes from these two species (*ShSULT-OR and SjSULT-OR*) to better understand the potential for resistance evolution will be critical if OXA derivatives are to be developed to control these parasites.

### 4. Implications of standing variation for drug resistance evolution

Standing variation in drug resistance genes has a major impact on how fast resistance evolves (2). If resistant alleles are already present in the population prior to treatment, drug resistance has the potential to spread much more rapidly because there is no waiting time for a resistance mutation to appear *de novo* (Fig. 7A) (3). An increasing number of examples support this view. The nematode *Caenorhabditis elegans* carries a large diversity of *β-tubulin* alleles allowing worms to be resistant to benzimidazoles (BZ) (32). In the filarial worm *Onchocerca volvulus*, standing variation appears to be involved in resistance to ivermectin (6). Standing variation is also involved in herbicide resistance in plants: resistance alleles from the weed *Alopecurus myosuroides* were identified in plant collection almost 100 years before herbicides were used (33). Antibiotic resistance provides particularly dramatic examples of standing variation: plasmids encoding multidrug resistance were found in frozen bacteria isolated from the permafrost demonstrating that antibiotic resistance plasmids were already present thousands of years ago (34, 35).

**Fig. 7.**
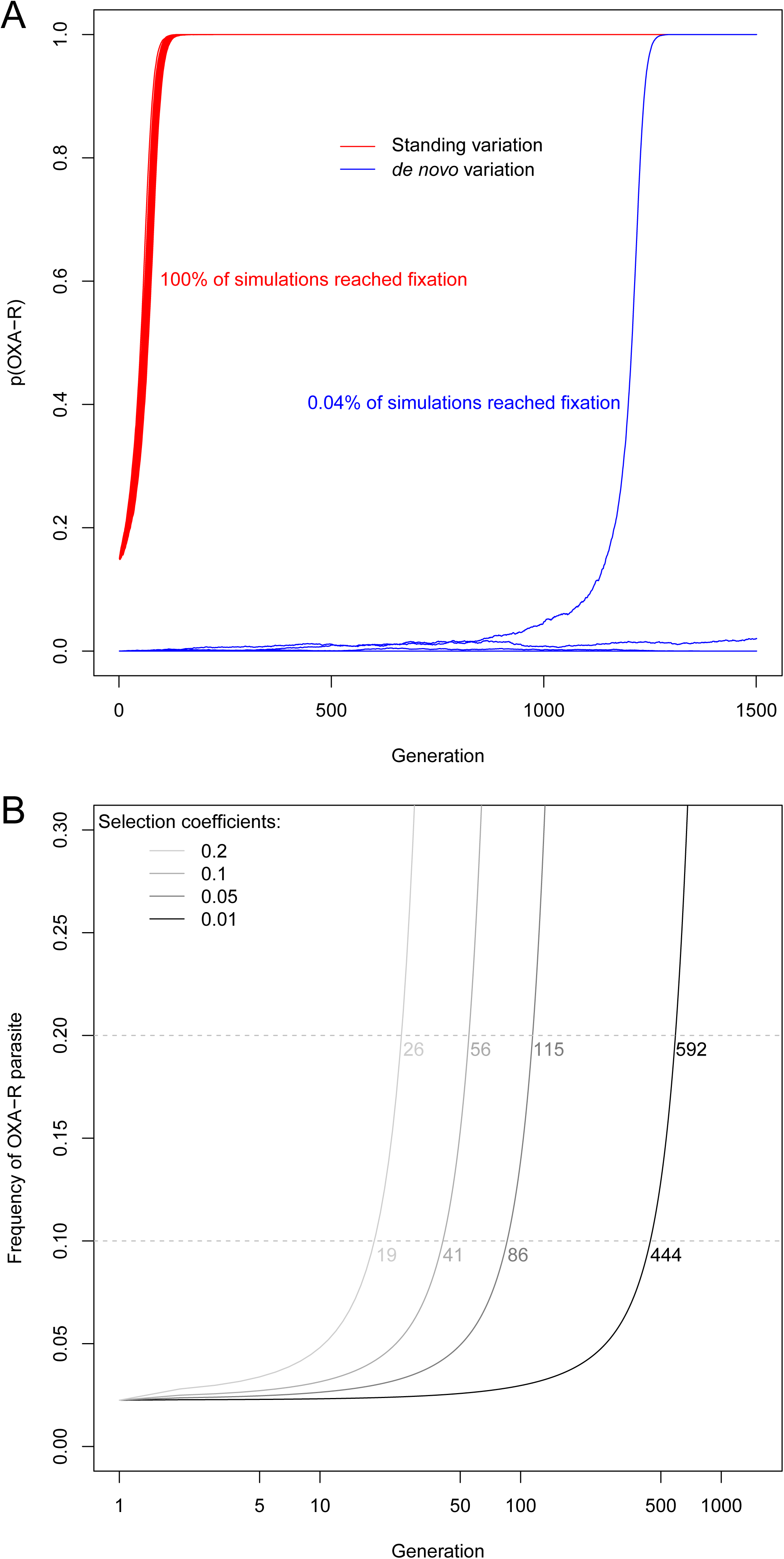
Impact of starting allele frequency on OXA-R allele change. (A) Monte Carlo simulation over 1,500 generations using strong selection (selection coefficient (s) = 0.1) on a population size (N) of 65,000 with a starting allele frequency (p(R)) of 0.15 for standing variation or 1/(2N) for new mutation. These simulations underestimate time to fixation for true *de novo* mutation, because we do not account for the waiting time for resistance to appear which is dependant on the rate of drug resistance mutations and N_e_(3). The starting frequency of 0.15 corresponds to the frequencies of OXA-resistance alleles observed in Kenya. (B) Change in frequency of resistant parasites (i.e. homozygotes for OXA-R alleles) from standing variation under a range of selection coefficients. The dashed line corresponds to the two thresholds (10% and 20%) at which treatment efficacy would be compromised. The numbers at the dashedlines correspond to the parasite generations needed to cross these thresholds. These predictions are deterministic, because we expect minimal stochastic variation when starting resistance allele frequencies are high.

We do not want to give the impression that standing variation is the only source of variation for evolution of drug resitance. In the case of other well studied helminth parasites, such as *Haemonchus contortus* and *Teladorsagia circumcincta*, resistance alleles are consistent with evolution through recurrent *de novo* mutations or very rare standing variants. While selection of rare pre-existing BZ resistance alleles was shown in French farms (36), selection of *de novo* resistance mutations, associated with different haplotypes, was shown to be common in UK farms (37).

The fact that there is strong evidence of standing variation in OXA resistance but not in BZ resistance may reflect the differences in the nature of the mutation underlying resistance. In the case of OXA, resistance is due to loss-of-function mutations resulting in non functional *SmSULT-OR*. As a consequence, mutations can occur in any position that disrupts substrate binding or protein structure. Hence mutations are expected to arise frequently and will exist in natural populations. In contrast, mutations underlying BZ resistance occur in specific position within the *β-tubulin* gene. As consequence, such mutations occur much more rarely. Furthermore because *β-tubulin* is an essential gene, mutations that disrupt protein structure and result in non functional proteins are strongly selected against and cannot reach high frequency.

Standing variation for OXA resistance alleles could arise by drift, or alternatively as a result of selective forces other than OXA treatment. For example, Hahnel *et al.* (32) demonsrated BZ resistance in natural population of C. elegans which are not targetted by BZ drugs and speculated that natural compound with similar structure to benzimidazoles could have pre-selected for benzimidazole resistance alleles. In the case of our OXA-R mutations in *SmSULT-OR*, we do not know what selective forces might be involved (see also discussion in section 5).

Drug resistance in parasitic helminths infecting livestock or humans is now recognized as a potential concern for control efforts, but was neglected for a long time using the argument that the slow reproduction rate of worms would greatly delay resistance evolution (38). The observation that resistance alleles exist as standing variation within multiple pathogen or parasite populations in the absence of drug treatment suggests previous optimism about the shelf life of drugs was poorly founded. If resistance alleles are already at relatively high frequency in populations when new drugs are deployed, resistance may evolve extremely rapidly (3, 5).

### 5. Prediction of the resistance spread

We can use the frequency of resistance allele to model resistance spread in parasite populations. In the case of OXA resistance, using an N_e_ estimate of 65,000 (39), OXA-R alleles spread from 15% starting frequency to fixation in 170 generations in 100% of simulations when selection is strong (s=0.1). Assuming a generation time between 3 months and 1 year, this is equivalent to a period of 42.5 to 170 years (Fig. 7A). However, if OXA-R alleles were evolving *de novo*, simulations indicate that fixation will occur rarely (0.04% of the simulations) and this would take 1,197 generations (corresponding to a period of 299.25 to 1,197 years) from the time a mutation arises. We can use the observed frequency of OXA-R alleles segregating in East African populations to make predictions about the spread of phenotypically resistant parasites (i.e. those homozygous for OXA-R alleles) (Fig 7B). If we use a frequency of 10% resistant parasites to denote unacceptable efficacy (this threshold is used for antimalarial drugs (40)) then this threshold will be reached in 19 to 444 generations (4.75 to 444 years) using a range of selection coefficients (from 0.01 to 0.2). The selection coefficients driving spread of drug resistance alleles in schistosomes is not known, but the range we have used covers measured selection coefficients driving drug resistance spread in another human parasite (*Plasmodium falciparum*) (41, 42). These simulations suggest that, while OXA-R allele frequencies are relatively high in East Africa, OXA derivatives could still be extremely effective for schistosome control in the short-medium term, particularly if deployed as a combination therapy, partnered with a second drug with a different mode of action (e.g. PZQ).

Our simple models for predicting spread of OXA-R alleles under drug selection ignore fitness costs of OXA-R alleles, which might limit the rate of spread. The fitness cost of inactive SmSULT-OR in the absence of drug treatment still needs to be assessed accurately (13) but the occurrence of natural homozygotes for defective alleles suggests limited costs. The exact role of this enzyme in schistosome biology is not known. Sulfotransferases transfer the sulfo group from a donor (usually PAPS) to the substrate, resulting in inactivating or solubilizing of the substrate (43). Substrates can be endogenous signal molecules (hormones, neurotransmitters) or exogenous chemicals. In the latter case, sulfotransferases play a role in detoxifying natural or synthetic toxins (43), but sometimes they do the opposite by activating pro-drugs as observed with the anti-tumor drug N-benzyl indolecarbinols (44) or OXA (10, 45). Because the same enzyme can sulfate both endogenous and exogenous compounds, it is often difficult to determine their exact biological role. In schistosomes, we speculate that *SmSULT-OR* could have roles in regulating endogenous molecules as well as detoxifying chemicals present in their environment (*i.e.* in the feces, water, snail or human blood, etc.).

### 6. Molecular markers for resistance surveillance

Molecular markers of OXA-R now allow efficient monitoring of the distribution of OXA-R alleles in schistosome populations. This approach is widely used for other parasites and pathogens such as malaria parasite (46) and HIV (47). For schistosome, collections such as SCAN which contain thousands of parasite samples from multiple locations (17) will be extremely valuable for this work: regular sampling in different endemic regions will help to update the resistance landscape. However, not all identified *SmSULT-OR* mutations will lead to resistance, so it is critical that molecular screening is paired with functional evaluation or computational prediction, as we have done here.

PZQ is the only drug currently available to treat schistosomiasis and effort in mass drug administration has recently been expanded 10 fold (48), with a target of administering 250 million treatments per year, increasing selective pressure for PZQ resistance. We suspect that standing variation for PZQ resistance is also likely, because this trait is easy to select in the laboratory. This has been done by several groups using independent laboratory populations, and resistance has spread within very few generations (49–54), strongly suggesting that resistance alleles were already present in these laboratory populations. Furthermore, the laboratory populations used for selection were isolated before PZQ was available (23, 55) and so were not exposed to drug selection. Molecular markers for monitoring the frequency of PZQ-R in natural parasite populations would be extremely valuable, as demonstrated by our work on OXA-R.

## MATERIALS AND METHODS

### 1. Ethics statement

#### a. African samples

Samples from Senegal and Niger were collected as part of the EU-CONTRAST project (56), a multidisciplinary alliance to optimize schistosomiasis control and transmission surveillance in sub-Saharan Africa, as detailed in Webster *et al.* (57) with ethical approval granted by ethical committees of the Ministry of Health (Dakar, Senegal), and the Niger National Ethical Committee (Niamey, Niger) with additional ethical approval obtained from the St Mary’s Hospital Local Ethics Research Committee, R&D office (part of the Imperial College Research Ethics Committee (ICREC; EC no. 03.36. R&D no. 03/SB/033E)), in combination with the ongoing CONTRAST and Schistosomiasis Control Initiative (SCI) activities.

For the Tanzanian samples collected as part of the Schistosomiasis Consortium for Operational Research and Evaluation (SCORE), ethical approvals were granted by the Imperial College Research Ethics Committee and the SCI Ethical approval (EC no. 03.36. R&D no. 03/SB/033E); the National Institute for Medical Research (NIMR, reference no. NIMR/HQ/R.8a/Vol. IX/1022); University of Georgia Institutional Review Boards, Athens, GA (2011-10353-1).

Following routine procedures in the field, the objectives of the study were first explained to the local village chiefs and political and religious authorities who gave their consent to conduct the study. Written consent for the schoolchildren to participate in longitudinal monitoring of the national control programme for schistosomiasis was given by head teachers, and/or village chiefs where there were no schools, due to the fact that in African villages, written consent of the child’s guardian is often very difficult to obtain owing to the associated impoverished conditions and often low literacy. Each individual child also gave verbal consent before −1 recruitment. Following sampling, a praziquantel treatment (40 mg.kg^−1^) was offered to infected participants.

#### b. Omani samples

We obtained ethical clearance from the Sultan Qaboos University and the Ministry of Health of Oman to use positive stool samples for schistosomiasis collected by the Ministry of Health of Oman during epidemiological screening. Ethical approval was given by the Medical Research and Ethical Committee (MREC) of the Non-Communicable Disease Control Section of the Directorate General of Health Affairs, Headquarters, Ministry of Health, Oman (no. MH/DGHA/DSDC/NCD/R&S/167/01).

We obtained ethical approvals of animal studies in France from the French Ministère de l’Éducation Nationale, de la Recherche et de la Technologie, from the French Ministère de l’Agriculture et de la Pêche (agreement no. A 66040), and from the French Direction Départementale de la Protection des Populations (no. C 66-136-01 with prefectoral order no. 2012-201-0008). Certificate for animal experimentation was given to H.M. (authorization no. C 66.11.01; articles no. R 214-87, R 214-122 and R 215-10). Housing, breeding and animal care followed the guidelines of the French CNRS. The different protocols used in this study had been approved by the French veterinary agency from the DRAAF Languedoc-Roussillon (Direction Régionale de l’Alimentation, de l’Agriculture et de la Forêt), Montpellier, France (authorization no. 007083).

### 2. Sampling

#### a. African samples

*S. mansoni* miracidia were collected from individual patients in three different West African countries in 2007: 27 patients from two locations in Senegal, and 17 patients from two locations in Niger (supp. table 3) (57, 58). Additionally, *S. mansoni* miracidia were collected as part of the SCORE program from 64 children in seven villages on the shores of Lake Victoria in Tanzania (East Africa) in January 2012 (supp. table 3) (59).

Collection of miracidia was performed as previously described (57, 60). Briefly, individual stool samples from positive patients were homogenized through a mesh and washed through with water. The content was then transferred in a Pitchford funnel assembly, washed with additional water, and the filtered homogenate was drained into a Petri dish. The Petri dish containing the homogenate was then left in bright ambient light (not direct sunlight) to allow hatching of miracidia. Miracidia were visualized under a dissecting microscope and individually captured in 3-5 µL of water using a micropipette. Miracidia were then pipetted individually onto Whatman FTA cards for DNA preservation. Cards were allowed to dry for 1 hour and then stored for future research (57,58,60).

#### b. Omani samples

The Omani schistosome samples were collected during 8 field trips, from 2001 to 2015, in different areas in Dhofar, Oman and originated from either *Homo sapiens, Rattus rattus* or *Biomphalaria pfeifferi* (supp. table 3): 11 patients from 4 localities, 2 rats from 2 localities, 26 snails from 3 localities.

We obtained schistosome adult worms from naturally infected rats and from laboratory infected mice. The mice were infected with cercariae from naturally infected snails or from laboratory snails infected with miracidia isolated from stool samples. We collected miracidia from stool samples as described in Moné *et al.* (61). We exposed *Biomphalaria pfeifferi* snails to miracidia as described in Mouahid *et al.* (62). We maintained laboratory and naturally infected snails at constant temperature (26°C) and balanced photoperiod (12h light / 12h dark) and fed them with fresh lettuce *ad libitum*. Prior to infection, we anaesthetized the mice by −1 intraperitoneal injection of an anesthetic solution (0.01 mL.g^−1^ of mouse body weight). We −1 prepared the anaesthetic solution using 0.5 mL of Rompun (20 mg.mL^−1^; Bayer) and 1 mL of −1 Imalgène (100 mg.mL^−1^; Rhône Mérieux) diluted in 8.5 mL of autoclaved NaCl 8.5 ‰. Abdomens of anesthetized mice were shaved and exposed during 1 hour to cercariae shed from laboratory or naturally infected snails. Cercariae from naturally infected snails were kept in 95° ethanol when the number of cercariae was too low for mouse infection. Laboratory infected mice or naturally infected rats were euthanized by intraperitoneal injection of sodium pentobarbital solution (1.1 mL diluted in 10 mL of ethanol 10%). We recovered adult worms by perfusion (63); the worms were washed in NaCl 8.5 ‰, and single worms (female or male) were fixed in 95° ethanol and preserved at −20°C. Worms and cercariae were washed in 1X TE buffer for 1 hour prior to DNA extraction.

### 3. DNA processing and library preparation

We amplified DNA and sequenced exomes from single miracidia preserved on FTA cards (1 miracidium per patient) for African samples or from DNA extracted from worms or cercariae for Omani samples. FTA-preserved samples were processed following our published protocol (64). Briefly we punched a 2 mm disc containing the miracidium from the FTA card, washed it with −1 the FTA Purification Reagent (GE Healthcare Life Sciences), rinsed it twice with TE buffer, and finally dried it. DNA from single worms and cercariae were extracted using the DNeasy Blood and Tissue kit (Qiagen) following the tissue protocol with an incubation time of 2h at 56°C and an elution in 200 µL. Samples were quantified using the Qubit dsDNA HS assay kit (Invitrogen).

We then performed whole genome amplification (WGA) on each FTA punch or on 2 µL or 4 µL of DNA solution using the Illustra GenomiPhi V2 DNA Amplification kit (GE Healthcare Life TM Sciences). Amplified DNA was purified with the SigmaSpin Sequencing reaction Clean-up (Sigma-Aldrich), following the manufacturer protocol. We quantified purified samples using the Qubit dsDNA BR assay (Invitrogen). DNA samples that failed WGA were cleaned up and concentrated using the Genomic DNA Clean & Concentrator kit (Zymo Research) following the manufacturer protocol and these samples with concentrated DNA were used to perform a new WGA.

Because WGA reactions are non-specific, we performed qPCR on each sample from FTA cards to quantify proportion of schistosome DNA (i.e., check for an excess of amplified DNA from the environment (water, fecal matter, etc.)). We excluded samples that showed less than 100 schistosome genomes copies in 20 ng of DNA which were not suitable for subsequent exome capture. We assessed genome copies by quantifyng the *S. mansoni α-tubulin* single copy gene (64).

We prepared exome capture libraries on the selected African samples and on all the successfully amplified Omani samples using the pre-capture pooling method of the SureSelect 2 XT Target Enrichment System (Agilent) (64). Finally, we sequenced libraries on a HiSeq 2500 and data were demultiplexed using the Casava pipeline. Raw sequencing data are accessible from the NCBI Sequence Read Archive under BioProject accession numbers PRJNA439266, PRJNA560069, PRJNA560070.

### 4. Alignment, variant calling and phasing

We aligned sequences as described in Le Clec’h *et al.* (64). Briefly, we aligned data against the v5 *S. mansoni* genome using BWA and SAMtools, realigned around indels using GATK, PCR duplicates were marked using picard and Q-score were recalibrated using GATK.

To identify variants in the *SmSULT-OR* (Smp_089320) gene, we performed a variant calling using FreeBayes (v1.1.0-46-g8d2b3a0) (65) in the first 3 Mb of the chromosome 6. We used a minimum base quality (-q) of 20 and a minimum mapping quality (-m) of 30. We included mutation sites identified previously (10, 13) in order to genotype these sites specifically for further comparison. We defined 4 populations: Caribbean (HR), West Africa (SN and NE), East Africa (TZ) and Middle-East (OM). We excluded variants supported by less than 4 reads using VCFtools (v0.1.14) (66). We then used vcf-subset from VCFtools to remove sites with reference alleles only. We adjusted variant positions using vt normalize (v0.5772-60f436c3) (67). We finally used Beagle (v4.1) (68) to first estimate genotypes and to then phase data. This approach resolves haplotypes using statistical models rather than by direct linkage of SNPs on short reads. Unphased variant calling data and scripts used to analyze the data are available on Zenodo (DOI: <u>10.5281/zenodo.2850876</u> and <u>10.5281/zenodo.3370039</u>, respectively).

### 5. Genetics analysis

We performed all the genetics analysis using phased exome data. We handled the VCF data in R (R v3.5.0 (69)) using the vcfR package (v1.8.0.9) (70). We determined the longest haplotype block from the samples carrying the p.E142del using a custom R code after filtering invariable sites in these samples. We built the haplotype network using the R package poppr (v2.8.1) (71, 72) after selecting variants within the longest haplotype coordinates showing no more than 20% of missing data. We computed the frequency of the OXA-R allele using a custom R script and functional annotation obtained through a custom bash script.

We generated haplotype sequences of the coding sequence only for the Old World samples using the updated phased VCF data exported from R and the BCFtools (v1.2) (73). We generated haplotypes of the South American samples from previous genotyping using a custom bash script. We included the *SmSULT-OR* homologue sequence from S. rodhaini previously identified (13). We aligned the haplotypes using Clustal Omega (v1.2.2) (74). We determined nucleotide diversity, number of synonymous and non-synonymous sites using DnaSP software (v6.12.01) (75). Populations were defined as described in the previous section. Scripts used to analyze the data are available on Zenodo (DOI: <u>10.5281/zenodo.3370039</u>).

### 6. Recombinant *SmSULT-OR* protein production and OXA activation assay

We produced recombinant *SmSULT-OR* proteins following Chevalier *et al.* (13). Briefly, mutations were introduced in the cloned *SmSULT-OR* gene sequence (Smp_089320; GenBank accession no. HE601629.1) and introduced into *Escherichia coli* for protein production. Proteins were extracted from bacterial culture and purified on an affinity chromatography column. His-tag was removed using Tobacco etch virus (TEV) protease, the solution was dialyzed overnight and passed through affinity chromatography again to remove His-tag and TEV protease. The sample was loaded onto a GE-pre-packed Q anion exchange column and eluted, and the pooled −1 fractions were dialyzed overnight. The protein was finally concentrated at 10 mg.mL^−1^.

We tested the impact of a subset of the mutations observed on OXA activation using an *in vitro* assay (13). We tested the ability of recombinant *SmSULT-OR* proteins to activate OXA in a protease inhibitor cocktail with sheared *S. mansoni* gDNA as a final target for the tritiated OXA pre-mixed with ATP, MgCl_2_ and PAPS co-factor. After 2.5 h of incubation at 37°C, the reaction was stopped and DNA was extracted three times. Radioactivity in the remaining aqueous phase containing the DNA and in water (blank) was counted in a liquid scintillation spectrometer. Blank values were subtracted from sample values. We performed three independent reactions for each recombinant protein.

### 7. Evaluation of impact of mutations on protein stability

We first visually assessed the scored mutations on the *SmSULT-OR* protein structure (PDB code 4MUB (10)) using the mutagenesis function of PyMol software (v2.3.0; Schrödinger, LLC).

We then used the Rosetta package (v3.9) to test the impact of scored mutations on protein stability. We used the ddg_monomer application to compute the difference in free enthalpy (*ΔΔG*) between the mutated (amino acid substitutions only) and the wild-type protein. The higher the *ΔΔG*, the more unstable is the mutated protein. For this, we prepared ligand files (OAQ and A3P) by downloading SDF files containing all ligand structures from *SmSULT-OR* crystals available from the Protein Data Bank website. We modified SDF files to add hydrogens using Openbabel (v2.4.0) (76), and generated params files using molfile_to_params.py from Rosetta. We used the structure of *SmSULT-OR* to generate a constraint file using the minimize_with_cst application and the convert_to_cst_file.sh script. We removed ligand information from the constraint file. For each mutation, we finally generated a resfile and ran the ddg_monomer application with 50 iterations and options adapted from the protocol 16 of Kellogg *et al.* (77). Scripts used to analyze the data are available on Zenodo (DOI: <u>10.5281/zenodo.3370039</u>).

### 8. Statistical analysis

We conducted statistical analyzes using R (v3.5.0) (69). We performed simulations using a modified version of the driftR simulator (78). The OXA activation assay data were tested for normality using a Shapiro-Wilk and compared by a parametric ANOVA followed by a Tukey post-hoc test.

## ACKNOWLEDGMENTS

This study was supported by NIH grants [R01-AI097576 (T.J.C.A.), R21-AI096277 (T.J.C.A.), R01-AI133749 (T.J.C.A.), R01-AI115691 and P50-AI098507 (P.T.L./P.J.H.)], World Health Organization [HQNTD1206356 (P.T.L.)], by the National Center for Advancing Translational Sciences (CTSA/IIMS award no. UL1-TR001120), the UTHSCSA Presidents Collaborative Research Fund (P.T.L./P.J.H.) and the Robert A. Welch Foundation [AQ-1399 (P.J.H.)]. The molecular work at Texas Biomed was conducted in facilities constructed with support from Research Facilities Improvement Program Grant (C06-RR013556) from the National Center for Research Resources (NIH). The AT&T Genomics Computing Center supercomputing facilities were supported by the AT&T Foundation and the National Center for Research Resources Grant (S10-RR029392). The X-ray Crystallography Core Laboratory is a part of the Institutional Research Cores at the University of Texas Health Science Center at San Antonio supported by the Office of the Vice President for Research and the Mays Cancer Center (NIH P30-CA054174). W.L was supported by a Cowles fellowship from Texas Biomedical Research Institute. CONTRAST was funded by the European Commission (FP6 STREP contract no. 032203). SCORE (https://score.uga.edu/) activities were funded by the University of Georgia Research Foundation Inc. (prime award no. 50816, sub awards RR374-053/4785416 and RR374-053/4785426), which is funded by the Bill & Melinda Gates Foundation for SCORE projects. SCAN is funded with support from the Wellcome Trust (grant no. 104958/Z/14/Z). For the samples collected as part of the EU-CONTRAST project we would like to thank Dr. Oumar T. Diaw and Dr. Moumoudane M. Seye (Institut Sénégalais de Recherches Agricoles, ISRA, route des Hydrocarbures, Bel Air, Dakar, Senegal) for their coordination of the field work and the collections in Senegal and Dr. Mariama Lamine for fieldwork coordination and the RISEAL team for collections in Niger. Also, we would like to thank Dr. Fiona Allan and Ms. Muriel Rabone for sample and data curation and preparation for material held in SCAN. For the collection of the Tanzania SCORE samples we would also like to acknowledge the hard work of Ms. Teckla Angelo for the fieldwork coordination, and Mr. Honest Nagai, Mr. Boniface Emmanuel, Mr. John Igogote, Dr. Sarah Buddenborg and Mr. Reuben Jonathan for collections in Tanzania. Dr. David Rollinson, Prof. Joanne Webster, Dr. Anouk Gouvras, Dr. Bonnie Webster and Dr. Aidan Emery are members of the London Centre for Neglected Tropical Disease Research, a collaboration between the London School of Hygiene & Tropical Medicine, the Natural History Museum, the Royal Veterinary College, Imperial College London and the Sanger Institute.

All the biological material from Oman was obtained thanks to the financial supports from the Ministry of Health in Oman, the Sultan Qaboos University (grant no. IG/MED/MICR/00/01), the French Ministry of Foreign Affairs (French Embassy in Oman) (grants nos. 402419B, 402415K and 339660F), the CNRS-Sciences de la Vie (grants no. 01N92/0745/1 and 02N60/1340), the CNRS-Direction des Relations internationales (grants no. 01N92/0745 and 02N60/1340, and the PICS-CNRS no. 06249 : FRANC-INCENSE), the University of Perpignan and the National Institute of Health (NIH) (grant no. R01-AI133749 T.J.C.A.; Subaward no. 53409 H.M.).

## SUPPLEMENTARY TABLES

Supp. table 1 – Library sequencing statistics.

Supp. table 2 – Mutations scored in samples from West Africa (Senegal and Niger), East Africa

(Tanzania) and Middle East (Oman).

Supp. table 3 – Sampling information for the schistosome material used.

## Notes

#### Summary of Updates

Revision after first peer review. Results and discussion amended, figures improved.

